# Pediatric High Grade Glioma Resources From the Children’s Brain Tumor Tissue Consortium (CBTTC) and Pediatric Brain Tumor Atlas (PBTA)

**DOI:** 10.1101/656587

**Authors:** Heba Ijaz, Mateusz Koptyra, Krutika S. Gaonkar, Jo Lynne Rokita, Valerie P. Baubet, Lamiya Tauhid, Yankun Zhu, Miguel Brown, Gonzalo Lopez, Bo Zhang, Sharon J. Diskin, Zalman Vaksman, Children’s Brain Tumor Tissue Consortium, Jennifer L. Mason, Elizabeth Appert, Jena Lilly, Rishi Lulla, Thomas De Raedt, Allison P. Heath, Alex Felmeister, Pichai Raman, Javad Nazarian, Maria Rita Santi, Phillip B. Storm, Adam Resnick, Angela J. Waanders, Kristina A. Cole

## Abstract

**Background:** Pediatric high grade glioma (pHGG) remains a fatal disease. Increased access to richly annotated biospecimens and patient derived tumor models will accelerate pHGG research and support translation of research discoveries. This work describes the pediatric high grade glioma set of the Children’s Brain Tumor Tissue Consortium (CBTTC) from the first release (October 2018) of the Pediatric Brain Tumor Atlas (PBTA).

**Methods:** pHGG tumors with associated clinical data and imaging were prospectively collected through the CBTTC and analyzed as the Pediatric Brain Tumor Atlas (PBTA) with processed genomic data deposited into PedcBioPortal for broad access and visualization. Matched tumor was cultured to create high grade glioma cell lines analyzed by targeted and WGS and RNA-seq. A tissue microarray (TMA) of primary pHGG tumors was also created.

**Results:** The pHGG set included 87 collection events (73 patients, 60% at diagnosis, median age of 9 yrs, 55% female, 46% hemispheric). Analysis of somatic mutations and copy number alterations of known glioma genes were of expected distribution (36% *H3.3*, 47% *TP53*, 24% *ATRX* and 7% *BRAF* V600E variants). A pHGG TMA (n=77), includes 36 (53%) patient tumors with matched sequencing. At least one established glioma cell line was generated from 23 patients (32%). Unique reagents include those derived from a *H3.3* G34R glioma and from tumors with mismatch repair deficiency.

**Conclusion:** The CBTTC and PBTA have created an openly available integrated resource of over 2,000 tumors, including a rich set of pHGG primary tumors, corresponding cell lines and archival fixed tissue to advance translational research for pHGG.

**IMPORTANCE OF STUDY:** High-grade gliomas (HGG) remain the leading cause of cancer death in children. Since molecularly heterogeneous, preclinical studies of pediatric HGG will be most informative if able to compare across groups. Given their relatively rarity, there are few readily available biospecimens and cellular models to inform preclinical laboratory and genomic translational research. Therefore, the aim of this CBTTC study was to highlight the panel of pediatric HGG cases whose primary tumors have undergone extensive genomic analysis, have clinical data, available imaging and additional biospecimens, including tumor, nucleic acids, cell lines and FFPE tissue on a tissue microarray (TMA).

## INTRODUCTION

Pediatric high grade glioma is a leading cause of pediatric cancer death, with most children dying of their disease within 2 years of diagnosis. Recent large genomic studies have shown that pHGGs arise along embryonic developmental lineages and are genomically and spatially distinct. ^1 2^ Anatomically midline tumors (e.g. thalamic, pontine) often have recurrent mutations in *H3.3* K27M (38% of all non-brainstem pHGG), and have earned the WHO classification of diffuse midline K27M mutant gliomas. In contrast, anatomically hemispheric tumors are more likely to have recurrent mutations in *H3.3* G34R/V, *IDH1* and *SETD2*. Additional recurrent mutations occur in *TP53, ATRX* and *BRAF*. Indeed, these discoveries are prompting the development of *H3.3* K27M targeted therapies and improved interpretation of high grade glioma clinical trial results. ^3 4^

In order to advance biological studies and clinical translation of these genomic discoveries in pHGG, it is necessary to 1) increase the availability of pre-clinical models linked to patient biospecimens as well as publicly available pHGG genomic datasets and 2) present the linked genomics data and clinical annotations in a manner that is accessible to the general scientific community. The Children’s Brain Tumor Tissue Consortium (CBTTC) is an international multi-institution biorepository that aims to integrate genomic and molecular research with biorepository management. Here we describe the generation and characterization of a highly annotated set of non-brainstem pediatric high grade glioma reagents freely available to the community to accelerate research of this devastating disease.

## MATERIALS AND METHODS

### General and laboratory methods

#### CBTTC biorepository query

The CBTTC (cbttc.org) is a collaborative, multi-institutional research program dedicated to the study of childhood brain tumors. The consortium currently includes 16 institutions worldwide (see Appendix). The CBTTC open source biorepository (https://eig.research.chop.edu/cbttc/) is available to member and non-member investigators through a submission request system. A research proposal request for biospecimens and/or data (clinical, imaging, histology slides, genomics) goes through a peer review process to approve specimen distribution and data sharing.^5^ After approval, the specimens are delivered to investigators while CBTTC data is available for viewing and download from the Gabriella Miller Kids First Data Resource Center (KF-DRC, https://kidsfirstdrc.org). Clinical reports can be downloaded as PDFs and Aperio SVS histology images viewed with QuPath. ^6^ Pre- and post-operative MRI images and reports are also available for a subset of the CBTTC subjects.

#### Processed Genomic Data

Next generation sequencing of over 1200 tumors from the CBTTC was completed in 2018 and deposited in the KF-DRC, as part of the Pediatric Brain Tumor Atlas (PBTA) as described below. Once approved, data is available for local download, analysis in the cloud-based data sharing platform CAVATICA or visualization of processed data in PedcBioPortal.^7^ For this study, CBTTC pHGG/PNET tumor sequences from the PBTA release 1 date of 10/30/18 were re-harmonized in 3/2019 and uploaded into PedcBioPortal (https://pedcbioportal.org), with proteomic data available for select cases. ^7^ Somatic mutations and copy number alterations in known pHGG and mismatch repair genes from PedcBioPortal were included if they were considered likely pathogenic by the OncoKB level of evidence tiers 1-4. ^8^

#### Human Subject Research Protections

All subjects were consented to tissue and data collection through CBTTC institutional IRB approved protocols. The KF-DRC is recognized as a National Institutes of Health (NIH) trusted partner for data sharing protections.

#### Identification of pHGG specimens for cell line generation

A search of the CBTTC biorepository (>3000 unique patients) in November 2017, revealed 37 non-brainstem pHGG tumors that had been stored in freezing media and available for tissue dissociation and cell culture generation as described below. As it is now recognized that up to 30% of tumors previously identified as Primitive Neuroectodermal Tumors (PNETs) are pHGGs, ^9^ PNETs were included in the search, but only included if a H3.3 G34R (n=2) or K27M (n=1) mutation was present.

#### Tissue dissociation and cell culture

All cell lines were generated by the CBTTC from prospectively collected tumor specimens stored in Recover Cell Culture Freezing media (Gibco, cat. 12648010) and dissociated as described in the Supplementary Methods. Up to three culture conditions were initiated based on the number of cells. For FBS cultures, a minimum density of 3×10^5^ cells/ml were plated in DMEM/F-12 medium supplemented with 20% fetal bovine serum (Hyclone, cat. SH30910.03), 1% GlutaMAX (Gibco, cat. 35050061), 100 U/mL penicillin-streptomycin and 0.2% Normocin (Invitrogen, cat. ant-nr-1). The remaining cells were plated in serum-free Neurobasal-A media (Gibco, 10888022) supplemented with 1% GlutaMAX, 1x B-27 supplement minus vitamin A (Gibco, cat. 12587010), 1x N-2 supplement (Gibco, cat. 17502001), 20 ng/ml epidermal growth factor (PeproTech, cat. AF-100-15B), 20 ng/ml basic fibroblast growth factor (PeproTech, cat. 100-18B), 10 ng/ml platelet-derived growth factor AA (100-13A), and 10 ng/ml platelet-derived growth factor BB (PeproTech, cat. 100-14B), 4μg/ml heparin (StemCell, cat. 07980), 100U/mL penicillin-streptomycin and 0.2% Normocin. Serum-free culture conditions included cells plated directly into media (SF) or on an extracellular Geltrex matrix denoted SF-ECM (Life Technologies, cat. A1413302) according to the manufacturer’s thin-gel method. Serum-free cells were initiated in culture with a minimum density of 1×10^6^ cells/ml. In cases where low cell counts prevented generation of a SF-ECM line at dissociation, established FBS cells were transitioned to the SF-ECM condition by the Cole laboratory staff at the lowest passage possible.

### Genomic Methods

#### DNA/RNA extraction, library preparation and sequencing

DNA extraction from 10-20 mg frozen tissue or 2×10^6 cells pellet was performed at Biorepository Core (BioRC) at Children’s Hospital of Philadelphia (CHOP). Briefly, tissue was lysed using a Qiagen TissueLyser II (Qiagen, USA) set at 2×30 sec at 18Hz using 5 mm steel beads. Qiagen AllPrep DNA/RNA/miRNA Universal kit including a CHCl3 extraction was utilized according to manufacturer’s protocol. DNA and RNA quantity and quality was assessed by PerkinElmer DropletQuant UV-VIS spectrophotometer (PerkinElmer, USA) and an Agilent 4200 TapeStation (Agilent, USA) for RINe and DINe (RNA Integrity Number equivalent and DNA Integrity Number equivalent). Library preparation and sequencing was performed by the NantHealth sequencing center. Briefly, DNA sequencing libraries were prepared for tumor and matched-normal DNA using the KAPA Hyper prep kit (Roche, USA); tumor RNA-Seq libraries were prepared using KAPA Stranded RNA-Seq with RiboErase kit. Whole genome sequencing (WGS) was performed at an average depth of coverage of 60X for tumor samples and 30X for germline. RNA samples were sequenced to an average of 200M reads. All samples were sequenced on the Illumina HiSeq platform (X/400) (Illumina, San Diego, USA) with 2 × 150bp read length. For the cell line sequencing, samples labelled with “CL-adh” correspond to the adherent FBS cell lines and those labelled “CL-susp” are the “S” serum free lines.

#### Variant calling and copy number alteration detection

The somatic variant and copy number variation detection workflow was setup using CWL (Common Workflow Language) on CAVATICA. The workflow uses BWA-MEM v0.7.17 ^10^ to align reads to the GRCh38 reference and SAMBLASTER v0.1.24 to mark duplicate reads, extract discordant/split reads ^11^, and sort the final BAM file. Base Quality Score Recalibration was applied using a model derived from known SNPs and InDels of HapMap, 1000 Genomes, dbSNP138, and Mills Gold Standard Calls databases. GATK4 v4.0.3.0 HaplotypeCaller was used to generate germline gVCF files, merge BAMs, and convert to CRAM as for both germline and tumor samples. Strelka2 v2.9.3 was applied on the aligned tumor-normal pairs to call point somatic mutations and short indels detection ^12^, Control-FREEC v8.70 was used to detect copy number alterations (CNAs) ^12^, and MANTA v1.4.0 was used for large structural variant detection. ^13^ The single nucleotide variants and indels were then annotated by SnpEff v4.3t and Variant-Effect-Predictor v93. ^14,15^ Only select mismatch repair (*MSH2, MSH6, PMS2, POLE*) and glioma-associated (*H3F3A, TP53, ATRX, BRAF, IDH1, NF1*) gene mutations that ‘PASSED’ Strelka2 filters and had predicted protein damaging effects were used for further analysis. For CNAs, both the Wilcoxon test and Kolmogorov-Smirnov test were performed and events < 10kb in length with p-value > 0.01 were filtered out.

Matched normal blood DNA was screened for rare (ExAC MAF <0.001) damaging mutations and focal deletions in a select panel of mismatch repair deficiency hereditary predisposition genes (*MSH2, MSH6, POLE, MLH1 and PMS2*). Only those determined to be Pathogenic or likely Pathogenic by InterVar ^16^ or previously reported ^17^ were used for further analysis.

#### Tumor mutation burden analysis

To obtain somatic variants in the coding region, only somatic variants overlapping a consensus exome region (https://github.com/AstraZeneca-NGS/reference_data) were used. The total base pair length of the exome bed was calculated to be 159697302 bp, which was then used to calculate the Tumor Mutation Burden (TMB) per sample as: (Total variants in coding region/length of exome bed (bp)*1000000.

#### RNA expression quantification

Sequencing data generation and read quality control checks were performed by the NantHealth sequencing center. RNAseq reads were aligned to hg38 reference and gene isoforms quantified using the Toil workflow (STAR v2.61d, RSEM v1.3.1). ^18^ FPKM (Fragments Per Kilobase of transcript per Million) and TPM (Transcripts Per Million) values were generated. FPKM values were uploaded to PedcBioPortal and we used the TPM values for visualizing the RNAseq data for all the CBTTC samples.

### Cell line - primary tumor comparison

#### Expression comparison

Primary tumor samples and matched cell line RNAseq expression data were filtered to retain genes that are expressed (> 1 TPM). We used 589 patient tumors as a reference set: 99 Ependymoma, 87 High-grade glioma/astrocytoma (WHO grade III/IV), 269 Low-grade glioma/astrocytoma (WHO grade I/II), and 134 Medulloblastoma tumors. We identified the 5000 most variable genes in pediatric tumors, then further restricted the analysis by filtering to genes expressed in both our primary and cell lines samples. This set of 3203 genes was log2 transformed for the reference set, cell lines, and primary tissue samples and pairwise spearman correlation was performed among disease type, cell lines, and their corresponding primary tissues.

#### Somatic Mutational Signature Analysis

The deconstructSigs R package v1.8.0 ^19^ was used to determine sample-level trinucleotide mutational matrices for all non-silent somatic mutations. Cosine similarities were then calculated for each of the 30 COSMIC signatures using the “genome” normalization mode for BSgenome.Hsapiens.UCSC.hg38.

## RESULTS

### The CBTTC cohort of pHGG primary tumors represent the spectrum of disease

The clinical co-variates, genomics and biospecimens of the pHGG set in this study are shown in Table 1, Table S1 and Figure 1. The cohort includes 81 pHGG and *H3.3* mutant PNET primary tumors with both WGS and RNA-seq from PedcBioPortal from 70 participants. An additional two cases (7316-1746 and 7316-599) with established cell lines and one case with proteomic data (7316-37) were included in the cohort. Therefore, the final set includes 73 participants representing 87 collection events from diagnosis and or progression/relapse. Tumors were from participants with a median age of 9 years old at diagnosis, with 97% of our cohort less than 18 years old at diagnosis. The cohort was balanced by sex as 55% of participants were female. The majority (60% of the tumors) were from initial or diagnostic specimens and nearly half (46%) were from a hemispheric location. There were two cases (7316-913, 7316-24) of secondary malignant glioma from patients treated with prior craniospinal radiation for medulloblastoma. Analysis of OncoKB somatic mutations and copy number alterations of a selected set of recurrently mutated pHGG genes were of an expected and representative distribution with 47% *TP53*, 36% *H3.3*, 24% *ATRX* and 7% *BRAF* V600E variants (Table S2 and Figure 1A).

**Figure 1.**
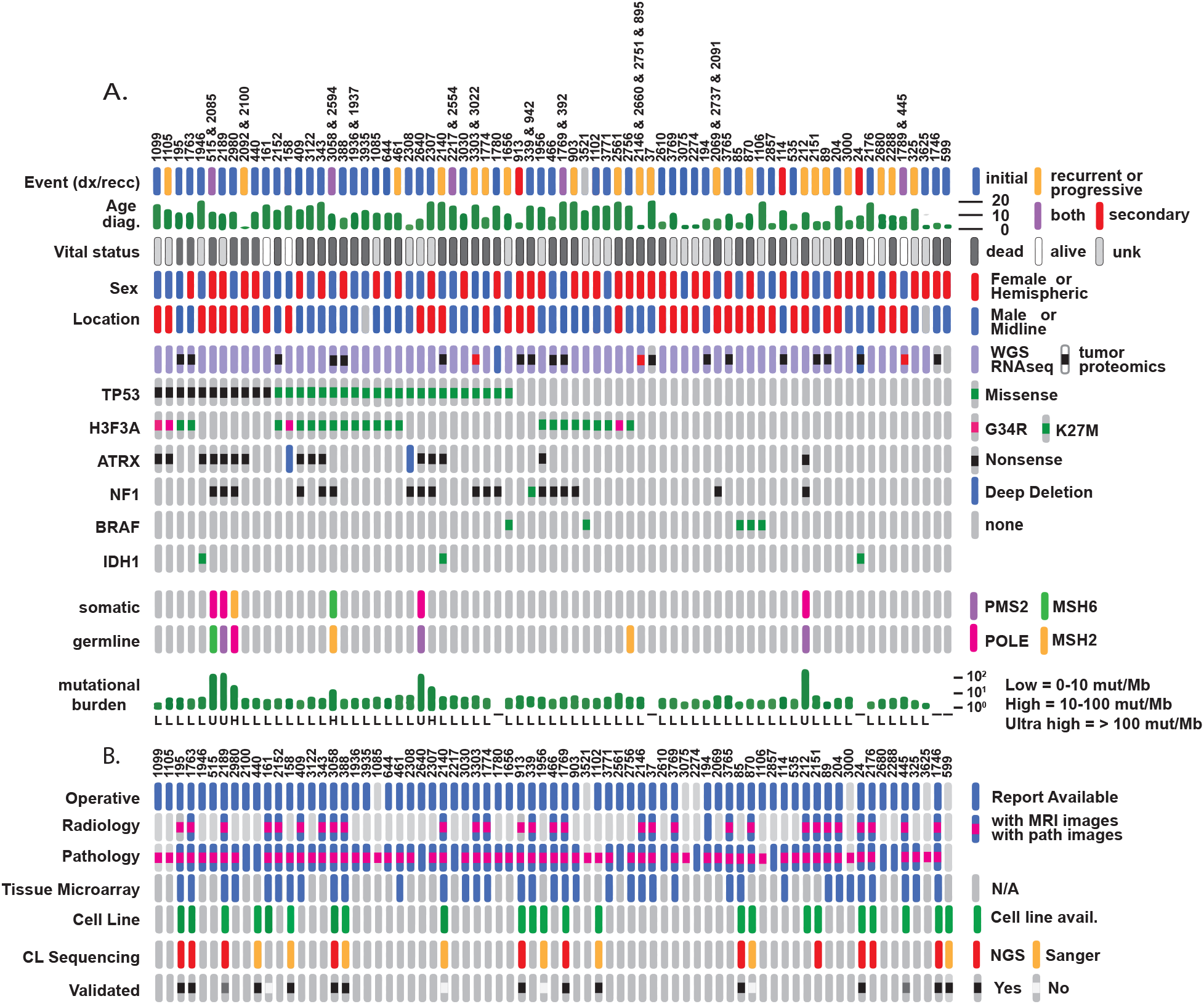
The CBTTC PBTA1 cohort and oncoprint. **A**. *Integrated Oncoprint* of the 73 participants in the complete pHGG cohort, with a corresponding key on the right. The demographic data such as event, age at diagnosis, sex and location were downloaded from CBTTC. Availability of WGS/RNAseq and tumor proteomics (red denotes both initial and recurrence) and select somatic pHGG genes (*TP53, H3F3A, ATRX, NF1, BRAF, IDH1, POLE, MSH2, MSH6*) as annotated by OncoKB were downloaded from PedcBioPortal or KF-DRC. Germline variants in *MSH2, MSH6, POLE and PMS2* and mutational burden were analyzed locally in CAVATICA. The bar graph of age and mutational on the oncoprint are approximate from Table S1 and S5 values. **B**. *Oncoprint of resources* available for the pHGG set with a descriptive key on the right. Availability of operative, radiology and pathology reports and imaging was downloaded from the KF-DRC manifest. Availability of pHGG FFPE samples on the TMA, cell line availability and sequencing were determined locally. Abbreviations: dx.: diagnosis; recc.: recurrence; NGS = next generation sequencing.

**Table 1.**
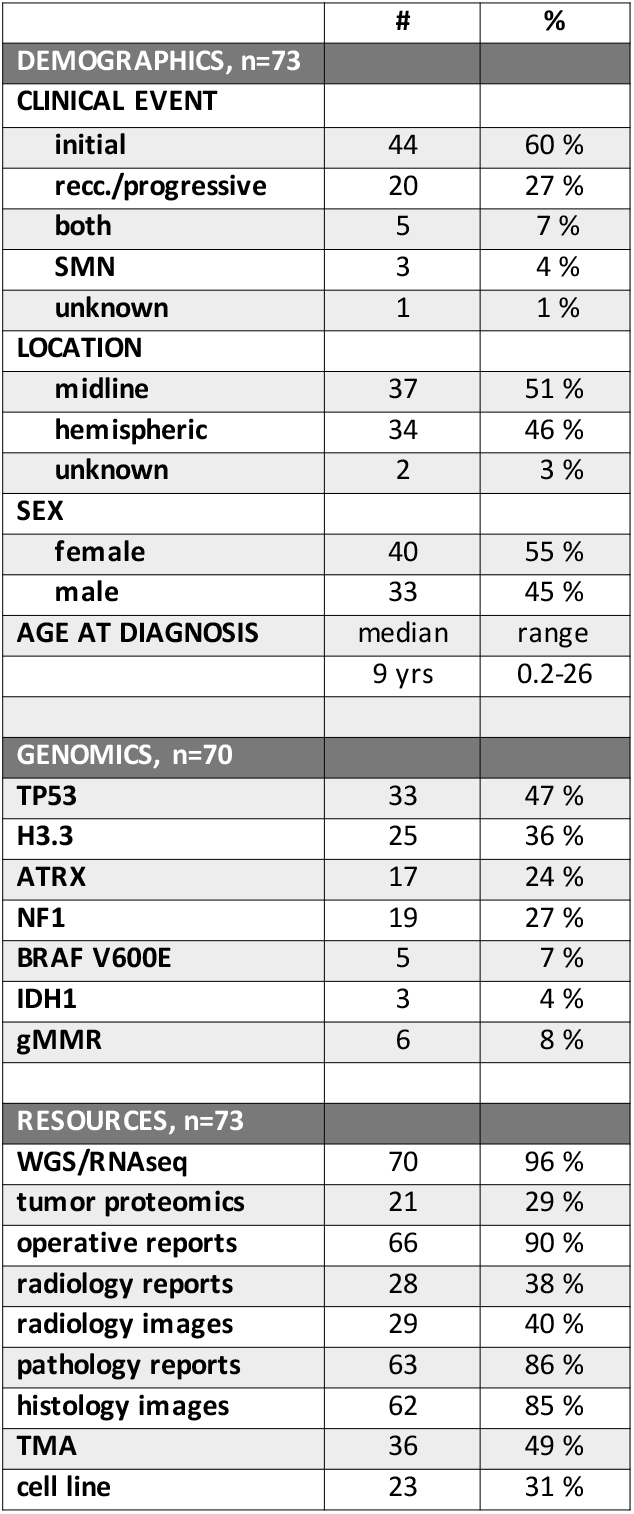
Summary of CBTTC pHGG cohort of the Pediatric Brain Tumor Atlas 1.

|  | # | % |
| --- | --- | --- |
| **DEMOGRAPHICS, n=73** |  |  |
| **CLINICAL EVENT** |  |  |
| initial | 44 | 60 % |
| recr./progressive | 20 | 27 % |
| both | 5 | 7 % |
| SMN | 3 | 4 % |
| unknown | 1 | 1 % |
| **LOCATION** |  |  |
| midline | 37 | 51 % |
| hemispheric | 34 | 46 % |
| unknown | 2 | 3 % |
| **SEX** |  |  |
| female | 40 | 55 % |
| male | 33 | 45 % |
| **AGE AT DIAGNOSIS** | median | range |
|  | 9 yrs | 0.2-26 |
| **GENOMICS, n=70** |  |  |
| TP53 | 33 | 47 % |
| H3.3 | 25 | 36 % |
| ATRX | 17 | 24 % |
| NF1 | 19 | 27 % |
| BRAF V600E | 5 | 7 % |
| IDH1 | 3 | 4 % |
| gMMR | 6 | 8 % |
| **RESOURCES, n=73** |  |  |
| WGS/RNAseq | 70 | 96 % |
| tumor proteomics | 21 | 29 % |
| operative reports | 66 | 90 % |
| radiology reports | 28 | 38 % |
| radiology images | 29 | 40 % |
| pathology reports | 63 | 86 % |
| histology images | 62 | 85 % |
| TMA | 36 | 49 % |
| cell line | 23 | 31 % |

As evident from the Oncoprint (Figure 1A) and as analyzed in PedcBioPortal, we were able to identify previously known co-occurring and mutually exclusive pHGG mutations. These include mutually exclusive *H3.3* G34 and K27M mutations^1^ and co-occurring mutations of *ATRX* with H3.3 G34R^20^ or NF1^21^. In this study, however, we also found that tumors with high confidence *PolE* mutations^22^ and ultra-hypermutation, also had significant cooccurrence of *ATRX* mutations (Bonferroni adj p value 0.013).

### CBTTC patient-derived pHGG biospecimens and resources are highly annotated and molecularly characterized

#### pHGG cell lines

The CBTTC biorepository had 37 pHGG tumor tissues stored in freezing media that were dissociated and cultured with four different culturing conditions, resulting in over 80 lines. At least one culture grew from 23/37 (62%) dissociation events from 23/73 (31%) patients in our cohort (Figure 1B). Information including prior patient therapy, culture and growing conditions including orthotopic xenografts, doubling times and validation status are described in Table 2 and Table S3. A subset of the cell lines from patients was sequenced by WGS and RNA-seq by the CBTTC or targeted Sanger sequencing where primary tumor variants could be identified (Figures 1B and 2 and Table S2). Of the 23 unique patient lines, pHGG driver mutations found in patient tumors validated in 13 derived cell lines, did not validate in four, and six pairs did not harbor known HGG mutations. For the 7316-445 cell line there was absent ATRX protein expression of the tumor on the pHGG TMA (not shown) and in the cell line by Western blot (Figure 3H). The 7316-2189 participant had a family history of childhood brain cancer and clinical diagnosis of Turcot Syndrome on the pathology report (Table S1). The 7316-2189 primary tumor and cell line both had low *PMS2* expression by FPKM on RNA sequencing and a previously described germline PMS2 mutation.^17^ The cell line panel represents the spectrum of pHGG genomics with eight unique patient cell lines with *H3.3* mutations (35%), six with *TP53* mutations (26%), one with a *BRAF* V600E mutation and one with a *KRAS* Q61H mutation. It is noteworthy that all cell lines matched their corresponding primary tumor identity by STR profiling, and also clustered with the cohort of 87 High-grade glioma/astrocytoma (WHO grade III/IV) by expression profiling (Figure 2A). Further studies will determine whether the cell lines that did not validate by sequencing represent non-tumor tissue or are sub-clones of the primary tumors.

**Figure 2.**
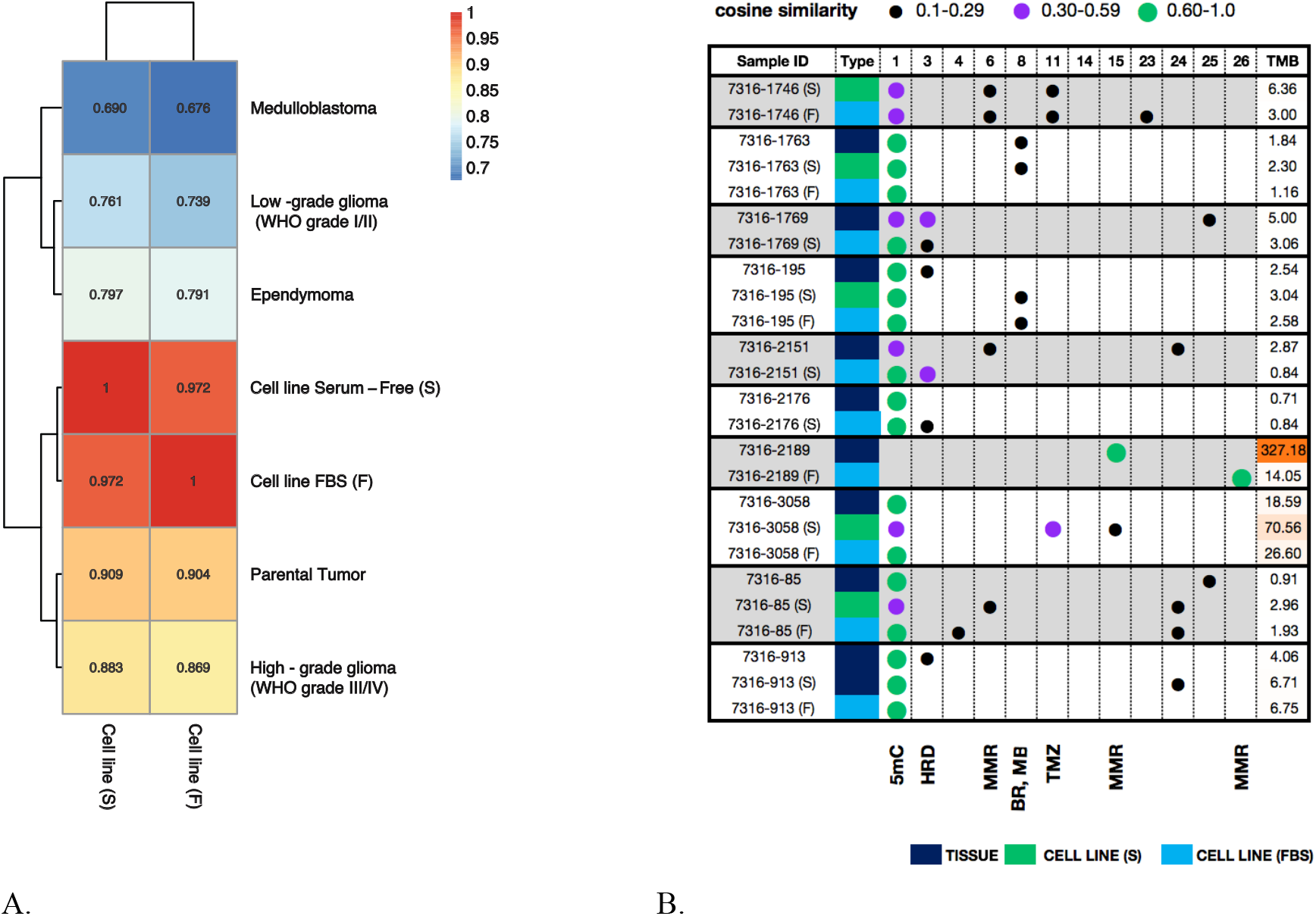
Parental primary tumor and cell line comparisons. **A**. Gene expression correlations of pHGG cell lines, parental tumors and brain tumors in the Pediatric Brain Tumor Atlas Set clusters the pHGG cell lines, parental primary tumors (n=9) and High-grade glioma (WHO grade III/IV) tumors (n=87). **B**. Mutational signatures of the pHGG cell lines (n=10) with corresponding primary tumor (n=9) WGS. “F” is Adherent-FBS. “S” is suspension, serum-free. Type refers to mutational signature nomenclature (COSMIC v2). TMB is the coding tumor mutational burden per Mb. 5mC is a deamination of 5-methylctyosine to thymine signature. HRD is homologous recombination defect associated signature. MMR is defective DNA mismatch repair associated signature. BR and MB are abbreviations for breast cancer and medulloblastoma, respectively. TMZ is a temozolomide signature.

**Table 2.**
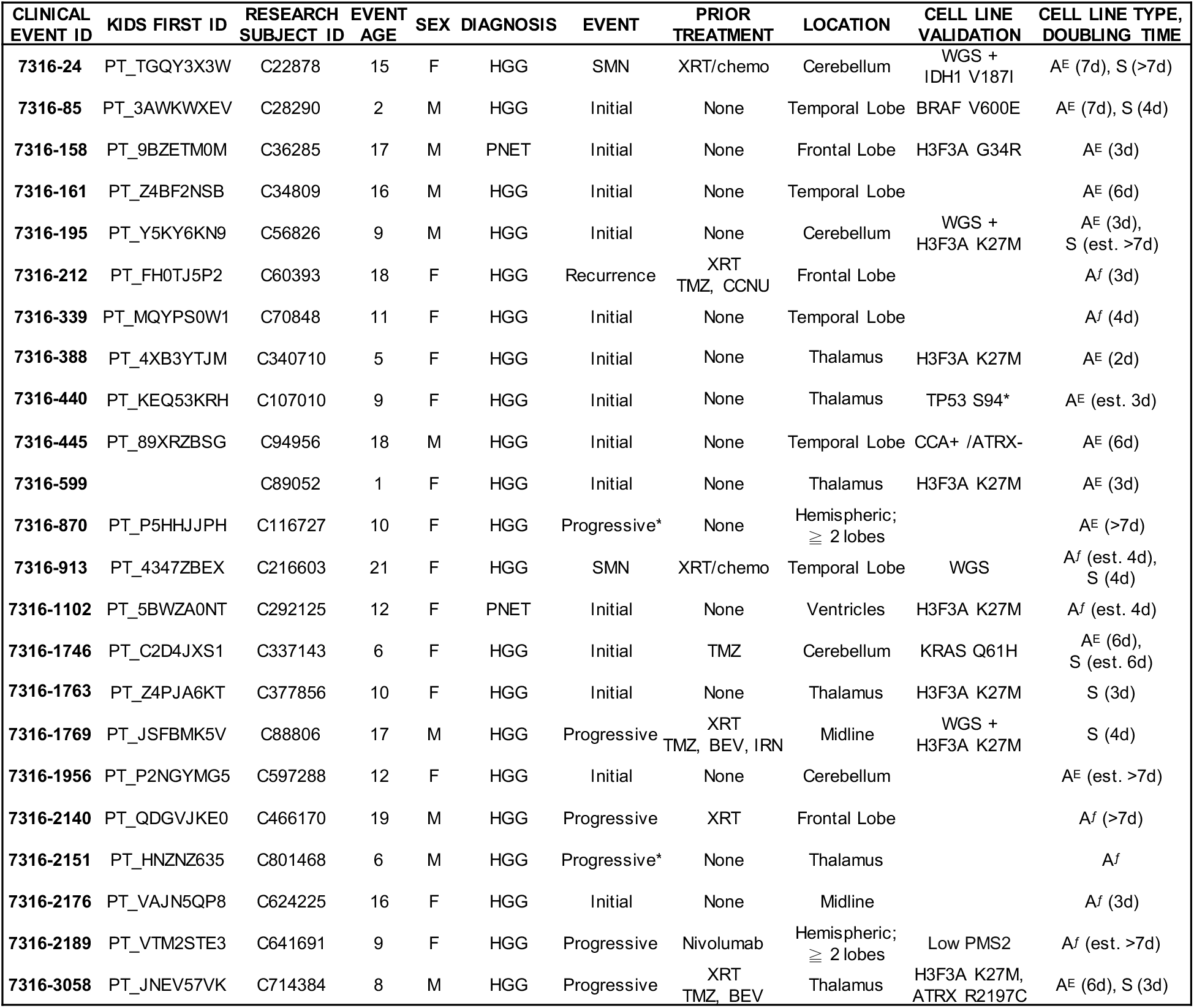
pHGG cell line table. Summary of established pHGG glioma patient derived cell lines. The clinical event ID is also called the “outside event ID” and is denoted by 7316-XXXX. It represents the base name of the cell line and the clinical event from which the line was established. The KidsFirst ID is the participant ID and has the prefix “PT_”. The Research ID is the CBTTC ID and starts with a “C”. Event age represents the age of the participant at the time of the event. Event designates whether the specimen was collected as an initial sample, at the time of progression or definitive surgery, at recurrence or was a second malignant neoplasm. The location summarizes the location of the tumor. The cell line validation column summarizes the gene and /or method of cell line validation. The cell line type is a list of the best line per sample designated as A (for adherent), S for suspension lines grown in serum free conditions, “f” for a line grown in fetal bovine serum, or “E” for an adherent line grown on an extracellular matrix in serum free conditions. The doubling times were determined by one of the three described methods. Details of the cell lines are found in Table S3.

Somatic mutations in cancer genomes may be caused by infidelity of the DNA replication machinery, exogenous or endogenous mutagen exposures, enzymatic modification of DNA, or defective DNA repair.^23^ To further characterize our set, we compared previously reported somatic mutational signatures of the cell lines to their parental tumors (Fig 2B and Table S5).^24^ All cell lines displayed the somatic mutational Signature 1 corresponding to spontaneous deamination of 5-methylcytosine (5mC), which is found in most cancer samples. Additional mutational signatures in the cell lines and parental tumors include those of homologous recombination deficiency (HRD) (Signature 3 in 7316-1769), prior alkylator therapy (TMZ) (Signature 11 in 7316-1746 and 7316-3058) and a double nucleotide substitution signature previously reported in breast cancer (BR) and medulloblastoma (MB) (Signature 8 in 7316-1763 and 7316-195). One of the most prevalent signatures were those for mismatch repair (MMR) deficiency (7316-2189) as described further below.

#### Additional CBTTC pHGG Resources

In addition to the cell lines, for each of the tumors in the CBTTC there are additional banked biospecimens including tumor tissue, tumor in freezing media, blood, plasma, CSF and both tumor and blood derived nucleic acids (Table S1). Of the pHGG cohort in this study, 36 patient tumors are also available on a tissue microarray (TMA) (Figure 1B), along with an additional 40 pHGG and control tissues. This valuable resource will enable researchers to examine the tumor microenvironment, discover cell surface immunotherapy target proteins or perform validation studies of the corresponding proteomic dataset. Finally, to facilitate integrative studies, most of the tumors in the pHGG set have corresponding redacted operative reports, pathology and radiology reports along with the relevant MRI images and histology slides (Figure 1B and Table S4).

### Use-case examples demonstrate the potential value of integrated resources to study pHGG

#### Resources to study hyper-mutation in pHGG

Pediatric high grade gliomas are one of the most common tumors in individuals with constitutional mismatch repair deficiency (CMMRD).^25^ They have germline loss of function of a mismatch repair gene (including *PMS2, MSH2, MLH1* or *MSH6*) and frequently acquire secondary somatic *POLE* or *POLD* mutations leading to ultra-hypermutant genomes with a high tumor mutational burden (TMB) > 100 mut/Mb. ^17,22^ It is these patients whose tumors have an exceptional clinical response to immune checkpoint inhibition ^26^ and require close tumor surveillance screening. ^27^ Individuals with germline *POLE* mutations often acquire secondary MMR somatic mutations and are hyper-mutant with 10-100 mut/Mb. ^17,22^ In our panel of pHGG, six patients (eight tumors) harbored germline mismatch repair mutations and hyper-mutation (Figure 1A) and one patient’s tumor is hyper-mutant without an identified germline variant (7316-2307). Four (7316-515 / 2085, 7316-2640, 7316-2189, 7316-212) had germline loss of function *PMS2* or *MSH6* mutations with high confidence somatic *POLE* mutations (Tables S2 and S6). ^22^ As predicted, these tumors were ultra-hypermutant with a TMB > 100 mut/Mb and had a somatic mutational Signature 15 (Figure 2B and Table S5) indicative of defective DNA mismatch repair. ^23^ One tumor (7316-2980) had a germline *PolE* mutation and secondary *MSH2* mutation and hyper-mutation (28 mut/Mb). Finally, three tumors (7316-2594/3058 and 7316-2756) were from two participants with heterozygous germline *MSH2* mutations (Lynch syndrome). Typically, individuals with Lynch syndrome do not develop cancer until adulthood. Additionally, the tumors from patient PT_JNEV57VK (7316-2594 / 7316-3058) were hyper-mutant. It is possible there were other genetic factors that contributed to glioma tumorigenesis at a young age, and therapeutic factors (i.e. radiation or alkylator therapy) that promoted tumor hyper-mutation.

Three of the hyper-mutant tumors were generated into cell lines (7316-2189, 7316-212 and 7316-3058). Interestingly, we defined the 7316-2189 tumor as ultra-hypermutant (TMB = 327 Mut/Mb), but its matched cell line was defined as hypermutant (TMB = 14.05 Mut/Mb). To further investigate, we ran somatic mutational signature analysis and found that the 7316-2189 tumor harbored a predominant signature 15 (cosine similarity = 0.91), supporting defective mismatch repair (MMR) in this tumor (Figure 2B). The predominant signature in the 7316-2189 matched cell line was signature 26 (cosine similarity = 0.91), an alternative trinucleotide context and signature of defective MMR, suggesting that even though the cell line is not ultra hypermutant, it still retains MMR deficiency.^23^

#### Availability of a H3.3 G34R mutant pHGG cell line

The development of preclinical cell and murine models of H3.3 K27M mutant midline glioma has significantly advanced the understanding of this histone biology in pHGG and also led to biomarker based clinical trials.^3^ However, the H3.3 G34R mutation is less common, occurring in 7% of pHGG and 16.2% of hemispheric tumors.^1^ Since there are fewer of these tumors, the models are also scarce, hindering discovery efforts for this subset of patients. In our pHGG set, there were two pHGG and two PNETs with a H3.3 G34R mutation, and as expected, were from adolescents with hemispheric tumors (Fig 1A). We were able to generate and characterize the 7316-158 H3.3 G34R line and examine additional resources to study this unique subset of pHGG. The resources include the 7316-158 MRI imaging (Figure 3A), diagnostic H&Es (Figure 3B) and immunohistochemistry slides. In addition, this tumor is on the pHGG TMA and shows protein loss of ATRX staining (Figure 3C). There are additional variants of interest as demonstrated on the tumor Circos plot (Figure 3D). The 7316-158 cell line (Figure 3E) was validated for the *H3.3* G34R mutation (Figure 3F) and absent ATRX protein (Figure 3H), and provides a resource for high throughput drug or genetic testing (Figure 3G, S1). Finally, the cell line from the matched recurrent specimen (7316-5317, Table S1) also has a validated *H3.3* G34R mutation. While the primary tumor sequence will be released in the upcoming updated PBTA2 release, the 7317-5317 cell line is currently available for distribution by the CBTTC.

**Figure 3.**
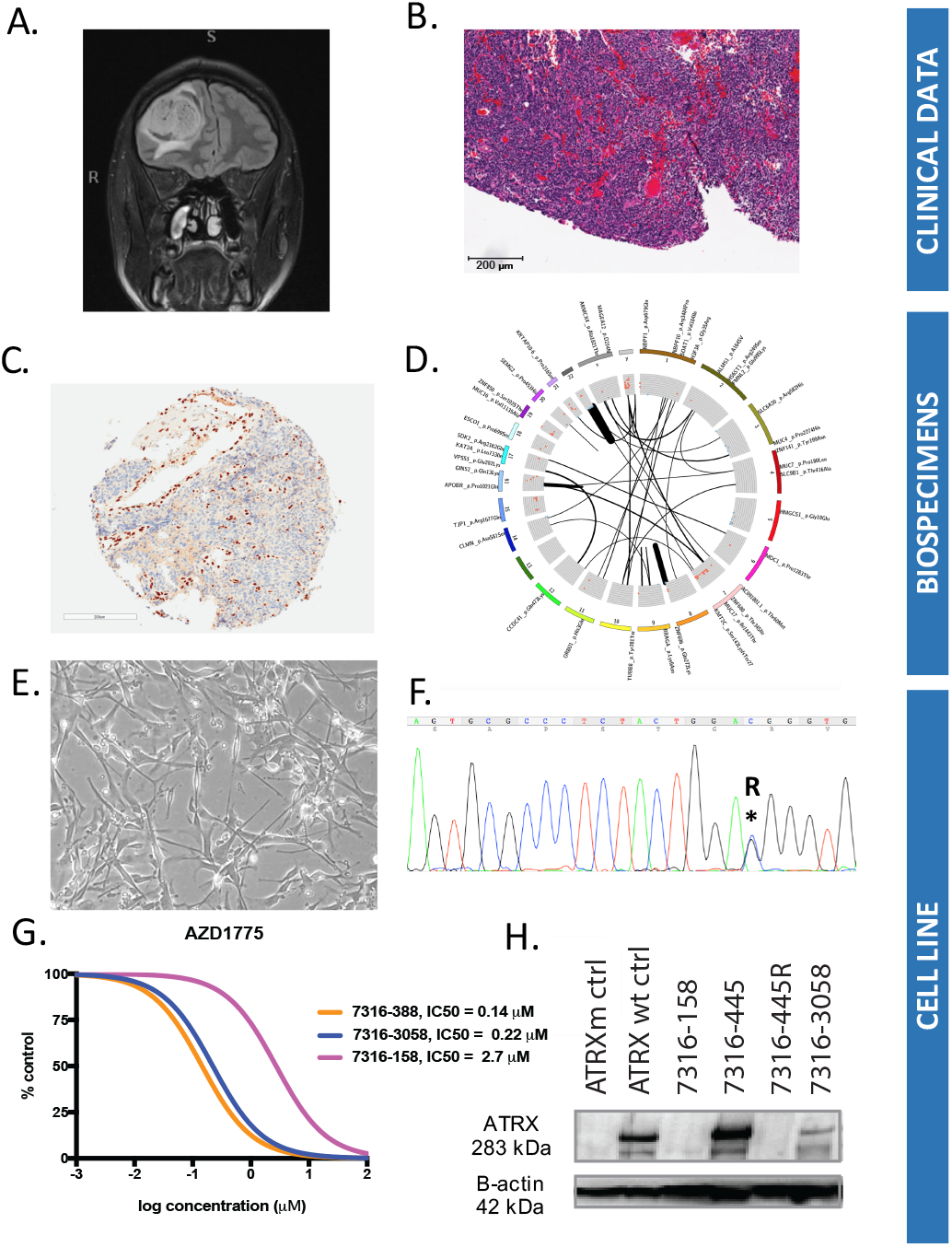
Integrated resources from participant PT_9BZETMOM (C36285) and the corresponding primary tumor (7316-158). **A.** Coronal MRI imaging demonstrates the pre-operative right frontal lobe tumor. **B.** Diagnostic H&E of the 7316-158 primary tumor. **C.** ATRX Immunohistochemistry of the 7316-158 tumor on the pHGG TMA shows absence of ATRX staining in the tumor cells (blue), but positive (brown) staining in cells of the microenvironment. **D.** Circos plot of the WGS of the 7316-158 primary tumor showing mutations (outer ring), copy number variation (middle ring) and structural variants (inner circle). **E.** Light microscopy of the 7316-158 cell line. **F.** Chromatogram of cell line DNA sequencing demonstrating the 7316-158 cell line H3.3 G34R mutation. **G.** Dose response curves of the 7316-158 (H3.3 G34R), 7316-3058 (H3.3 K27M) and 7316-388 (H3.3 K27M) cell lines treated with the Wee1 inhibitor AZD1775. **H.** ATRX Western Blot demonstrating loss of ATRX protein in the 7316-158 and control cell lines.

## DISCUSSION

Pediatric high grade gliomas remain incurable, emphasizing the need for collaborative efforts to openly share data and biospecimens to advance the care for children and adolescents with this rare disease. For example, integration of international pHGG genomic datasets in PedcBioPortal from the CBTTC (this study), the Herby Trial, Pacific Neuro-Oncology (PNOC) 003 clinical trial and International HGG consortium results in over 1500 cases of pHGG and diffuse intrinsic pontine glioma (DIPG).^1,28,29^ These datasets are expected to increase, lending statistical power to analysis of subsets of pHGG to advance discoveries and collaborations. For example, the second release of data for the CBTTC/PBTA is expected in 2019 and will include additional paired diagnosis and relapse specimens. Moreover, because the PedcBioPortal platform integrates with other non-brain tumor pediatric and adult cancer genomic datasets, the statistical power of discovery is further augmented. The same concept applies to pHGG models including cellular cultures, patient derived xenografts or murine models.^30^ Each has their unique advantages and limitations, but integration is essential to drive the field forward and necessary for cross-validation studies.

A unique aspect to the CBTTC is the prospective nature of the data collection and the open availability of data and biospecimens for follow up validation or additional discovery efforts. The CBTTC expects that the data and reagents generated from CBTTC specimens are returned to the scientific community as it is recognized that there is greater knowledge that can be learned from collaborative integrative analysis of tumors within a set. Finally, the need to share data and work collaboratively is not only scientifically rational, but is also mandated by the patients and parents who have supported this effort through donation of specimens, data and philanthropy.

This is the first report that focuses on the pHGG data and biospecimens that are available from the CBTTC and PBTA. The two use-cases describe patient cohorts that comprise subtypes of pHGG. We highlight the hypermutant pHGG cases and also *H3.3* G34R 7316-158 cell line, which is one of the first published available lines of its kind. Additional subsets include 1) specimens from multiple events which could evaluate resistance or response to therapy (n =11 patients), 2) secondary malignant glioma (n=3) and 3) analysis of tumor with recurrent mutations such as *NF1* or *ATRX*. Additional analyses of the set could include evaluation of the noncoding genome, structural variant analyses and interrogation of germline variation. There are also likely unique opportunities for neuro-radiologists and computational biologists to examine imaging characteristics of these highly characterized tumors.

There are several future initiatives that will enhance the existing pHGG data set and future CBTTC / PBTA studies. The CBTTC clinical data subcommittee recently completed a large data review and bioinformatics effort to provide clinical outcome data, including survival statistics for the CBTTC cohort to be available on PedcBioPortal. There are several current and prospective partnerships with the CBTTC to advance research and therapy for children with brain tumors. For example, a cohort tumors presented in this manuscript is included in the study of multi-omic single cell analyses as part of the NIH Cancer Moonshot Human Tumor Atlas Network (HTAN) and also Project HOPE. In addition, the CBTTC will continue its strong partnership with the Pacific Neuro-Oncology Consortium (PNOC) which is a multi-institution clinical trials network that facilitates rapid translation between basic, translational and clinical scientists to rapidly advance treatments for children with malignant brain tumors, including pediatric highgrade glioma.

## ACKNOWLEDGEMENTS

Clare Choi and Tasso Karras for technical support, Dan Martinez from the CHOP Pathology Core, Martha H. Williams, Tiffany Smith and Katie Boucher for assistance with cell line generation

## AUTHOR GROUPS & CONSORTIA

### CBTTC Principal investigators and Co-Investigators, by institution

1. Children’s Hospital of Philadelphia
2. Seattle Children’s
3. UPMC Children’s Hospital of Pittsburgh
4. Ann & Robert H. Lurie Children’s Hospital of Chicago
5. Benioff Children’s Hospital (UCSF)
6. Stanford University/Lucile Packard Children’s Hospital
7. Meyer Children’s Hospital Florence Italy
8. Weill Cornell Medicine Pediatric Brain and Spine Center
9. Children’s National Health System
10. Joseph M. Sanzari Children’s Hospital at Hackensack University Medical Center
11. Wake Forest Baptist Health
12. University of California Santa Cruz – Treehouse Childhood Cancer Initiative
13. The Beijing Tiantan Hospital Neurosurgery Center (Beijing, China)
14. Genebank (Beijing Genomics Institute – Shenzhen, China)
15. Dayton Children’s Hospital
16. The Hudson Institute of Medical Research (Melbourne, Australia)

### CBTTC Foundation Partners

Alan Stallings Fund, Alex Munoz Foundation, Andrew Noten Hudson Foundation, At Least Kids Foundation, Avery Lubrecht Foundation, BethAnn Telford, TeamBT, Brain Tumor Avengers, Brendan Bovard Fund for Brain Tumor Research, Bryce Hansen Bridge of Hope Fund, Charles and Pat Genuardi, Chester County Community Foundation, Children’s Brain Tumor Foundation and the Licensing and Merchandiser’s Association, Christopher Brandle Joy of Life Foundation, David and Deborah Calvaresi, Derrick and Lauren Roamelle, Dragon Master, Foundation, Eaise Family Foundation, Inc., Grayson Saves Foundation, Hanna Duffy Foundation, James and Nancy Minnick, Joseph T. Lentz Pediatric Brain Cancer Research Fund, Kayla’s Hope for Kids Fund, Kyle Daniel Kerpan Foundation, Lauren’s First and Goal, Miriam’s Kids Research Foundation, Naya Foundation, Pearce Q. Foundation, Inc., PLGA Foundation, Smiles for Jake Foundation, Stanley’s Dream, Suzanne Gilligan – Luke Forward, Swifty Foundation, The Christopher Court Foundation, The CJR Memorial Foundation, The Kortney Rose Foundation, The Lilabean Foundation, The Matthew Renk Foundation, The Timothy Pauxtis Foundation, Thea’s Star of Hope, Vs Cancer Foundation, Why Not Me? Foundation, Wylie’s Day Foundation

